# 16S rRNA phylogeny and clustering is not a reliable proxy for genome-based taxonomy in *Streptomyces*

**DOI:** 10.1101/2023.08.15.553377

**Authors:** Angelika B Kiepas, Paul A Hoskisson, Leighton Pritchard

## Abstract

Although *Streptomyces* is one of the most extensively studied genera of bacteria, their taxonomy remains contested and is suspected to contain significant species-level misclassification. Resolving the classification of *Streptomyces* would benefit many areas of study and applied microbiology that rely heavily on having an accurate ground truth classification of similar and dissimilar organisms, including comparative genomics-based searches for novel antimicrobials in the fight against the ongoing antimicrobial resistance (AMR) crisis. To attempt a resolution, we investigate taxonomic conflicts between 16S rRNA and whole genome classifications using all available 48,981 full-length 16S rRNA *Streptomyces* sequences from the combined SILVA, Greengenes, Ribosomal Database Project (RDP) and NCBI (National Center for Biotechnology Information) databases, and 2,276 publicly available *Streptomyces* genome assemblies. We construct a 16S gene tree for 14,239 distinct *Streptomyces* 16S rRNA sequences, identifying three major lineages of *Streptomyces*, and find that existing taxonomic classifications are inconsistent with the tree topology. We also use these data to delineate 16S and whole genome landscapes for *Streptomyces*, finding that 16S and whole-genome classifications of *Streptomyces* strains are frequently in disagreement, and in particular that 16S zero-radius Operational Taxonomic Units (zOTUs) are often inconsistent with Average Nucleotide Identity (ANI)-based taxonomy. Our results strongly imply that 16S rRNA sequence data does not map to taxonomy sufficiently well to delineate *Streptomyces* species reliably, and we propose that alternative markers should instead be adopted by the community for classification and metabarcoding. As much of current *Streptomyces* taxonomy has been determined or supported by historical 16S sequence data and may in parts be in error, we also propose that reclassification of the genus by alternative approaches is required.

**Impact Statement:** Accurate classification of microbes, usually in the form of taxonomic assignments, provides a fundamental ground truth or reference point for many aspects of applied microbiology including comparative genomics, identification of strains for natural product discovery, and dereplication of strains. Bacteria belonging to the genus *Streptomyces* are an important source of bioactive metabolites and enzymes in biotechnology, and proper understanding of their phylogeny aids understanding of the evolution of industrially important gene products and metabolites, and prioritization of strains for industrial exploitation. Taxonomic classification in the genus *Streptomyces* is complex and contested, and there are clear conflicts between taxonomies inferred from 16S rRNA and from whole genome sequences. Despite this, 16S sequence-based classifications are still widely used to infer taxonomic identity, to determine community composition, and to prioritise strains for study. We investigate a diverse and comprehensive set of *Streptomyces* genomes using whole-genome Average Nucleotide Identity (ANI) and 16S sequence analysis to delineate and compare classifications made using these approaches. We outline the genomic and 16S sequence landscape of *Streptomyces*, demonstrating that (i) distinct taxonomic species may share identical full-length 16S sequences, and (ii) in some instances, isolates representing the same taxonomic species do not share any common 16S rRNA sequence. Our results strongly imply that 16S rRNA sequence variation does not map to taxonomy sufficiently well to delineate *Streptomyces* species reliably, and that alternative markers should instead be adopted by the community. Much of current *Streptomyces* taxonomy has been determined or supported by historical 16S sequence data, and we therefore propose that reclassification within this group by alternative approaches is required.

**Data summary:** All code, raw and supporting data are publicly available from GitHub (https://github.com/kiepczi/Kiepas_et_al_2023_16S) and Zenodo (https://doi.org/10.5281/zenodo.8223787). The flowchart provided in `Supplementary File 28` provides an overview of analysis steps and serves as a guide through Supplementary Files generated during reconstruction of the 16S phylogeny. The flowchart in `Supplementary File 29` outlines the workflow processes and supplementary materials used for analysis of 16S rRNA sequences from *Streptomyces* genomes.

**Supplementary Data:** **Supplementary File 1: Generate figures using Python and R.** ZIP file containing all data, Python and R scripts to generate figures for this manuscript. (ZIP 40.9MB)

**Supplementary File 2: Raw 16S rRNA public databases.** Zip file containing four separate txt files with sequence IDs for public 16S rRNA databases used in this manuscript, and an additional txt file with Greengenes sequence taxonomy information, and a Python script used to map taxonomy information to sequences found in Greengenes v13.5. (ZIP 34.8MB)

**Supplementary File 3: Filtration of 16S rRNA public databases**. Zip file containing Python script used for filtration of the raw databases, and generated outputs. (ZIP 7.2MB)

**Supplementary File 4: Cleaning of the filtrated 16S rRNA local.** Zip file containing all bash and Python scripts used to clean the local full-length 16S rRNA local databases by removing redundant and poor quality 16S rRNA sequences. (ZIP 9MB)

**Supplementary File 5: Sequence Clustering.** Zip file containing a bash script used to cluster the full-length cleaned local 16S rRNA S*treptomyces* local databases at various thresholds, and provides txt files with accessions for representative sequences, and cluster members for each clustering threshold. (ZIP 40.8MB)

**Supplementary File 6: Analysis of taxonomic composition for each clustering threshold.** Zip file containing Python scripts, NCBI taxonomy input and all outputs generated used to determine the taxonomic composition for each clustering threshold. (ZIP 49.6)

**Supplementary File 7. Cluster sizes.** Empirical cumulative frequency plot showing cluster sizes generated for all clustering thresholds. (PDF 44KB)

**Supplementary File 8. Cluster taxID abundance.** Empirical cumulative frequency plot for unique numbers of taxID present at all clustering thresholds. (PDF 9KB)

**Supplementary File 9. MSA.** Zip file containing all Python and bash scripts, and additional data needed to generate and clean MSA for phylogenetic analysis. (ZIP 4.2MB)

**Supplementary File 10. Phylogenetic reconstruction.** ZIP file containing bash scripts used for phylogenetic reconstruction, and all generated outputs and log files. (ZIP 16.8MB).

**Supplementary File 11. Collapse branches.** ZIP file containing jupyter notebook used for collapsing branches with the same species names, and the collapsed tree in newick format. (ZIP 385KB)

**Supplementary File 12. Phylogenetic tree.** PDF file showing collapsed phylogenetic tree with marked branches with transfer bootstrap expectation support of >= 50%. (PDF 224KB)

**Supplementary File 13. Phylogenetic tree.** PDF file showing collapsed phylogenetic tree showing distribution of *Streptomyces albus* and *Streptomyces griseus.* (PDF 229KB)

**Supplementary File 14. Phylogenetic tree.** PDF file showing collapsed phylogenetic tree showing distribution of *Streptomyces albulus, Streptomyces lydicus* and *Streptomyces venezuelae.* (PDF 228KB)

**Supplementary File 15. Phylogenetic tree.** PDF file showing collapsed phylogenetic tree showing distribution of *Streptomyces clavuligerus* and *Streptomyces coelicolor.* (PDF 227KB)

**Supplementary File 16. Phylogenetic tree.** PDF file showing collapsed phylogenetic tree showing distribution of *Streptomyces lavendulae, Streptomyces rimosus* and *Streptomyces scabiei.* (PDF 228KB)

**Supplementary File 17. *Streptomyces* genomes.** Zip file containing bash scripts used to download *Streptomyces* genomes, and Python scripts used to check assembly status. The ZIP file also contains two separate txt files with *Streptomyces* genomes used in this manuscript: one file with all initial candidates, and a second file with replaced genomes. (ZIP 2.6MB)

**Supplementary File 18**. **Extraction of full-length and ambiguity free 16S rRNA sequences from *Streptomyces* genomes.** Zip file containing all Python and bash scripts used to extract full-length sequences from the filtered *Streptomyces* genomes. A single FASTA file with all extracted 16S rRNA sequences, and a single FASTA file with filtered sequences. A txt file with accession of genomes retained in the analysis. (ZIP 742KB)

**Supplementary File 19. ANI analysis among *Streptomyces* genomes with identical 16S rRNA sequences.** ZIP file containing all Bash and Python scripts used to determine taxonomic boundaries among *Streptomyces* genomes sharing identical full-length 16S rRNA sequences. All output and pyANI log files. (ZIP 37.1MB)

**Supplementary File 20. Network analysis of genomes based on shared 16S sequences.** ZIP file containing jupyter notebook with NetworkX analysis and all associated output files including. bash script for pyANI analysis runs on each connected component and all associated matrices, heatmaps and log files. (ZIP 29.3MB)

**Supplementary File 21. Interactive network graph.** HTML file containing interactive network graph of genomes sharing common full-length 16S sequences with each node colour corresponding to the number of connections/degrees. (HTML 4.7MB)

**Supplementary File 22. Interactive network graph.** HTML file containing interactive network graph of genomes sharing common full-length 16S sequences showing clique (blue) and non-clique (green) components. (HTML 4.7MB)

**Supplementary File 23. Interactive network graph.** HTML file containing interactive network graph of genomes sharing common full-length 16S sequences showing number of unique genera within each connected component. Each candidate genus is represented as a single node colour within a connected component. (HTML 4.7MB)

**Supplementary File 24. Interactive network graph.** HTML file containing interactive network graph of genomes sharing common full-length 16S rRNA sequences showing number of unique species within each connected component. Each candidate species is represented as a single node colour within a connected component. (HTML 4.7MB)

**Supplementary File 25 Interactive network graph.** HTML file containing interactive network graph of genomes sharing common full-length 16S rRNA sequences showing number of unique NCBI names within each connected component. Each NCBI assigned name is represented as a single node colour within a connected component. Gray nodes represent genomes currently lacking assigned species names. (HTML 4.7MB)

**Supplementary file 26.** Intragenomic 16S rRNA heterogeneity within 1,369 Streptomyces genomes which exclusively contain only full-length and ambiguity symbol-free 16S rRNA sequences. A total of 811 genomes containing single 16S rRNA sequences are not shown. (PDF 8KB)

**Supplementary File 27. Distribution of 16S copies per genome with a distinction between unique and total copies for genomes at assembly level complete and chromosome.** (PDF 7KB)

**Supplementary File 28. Schematic workflow for construction of the full-length 16S rRNA *Streptomyces* phylogeny**. Each arrow represents a process and is annotated with script used and corresponding supplementary file. Output/data files, and the number of remaining sequences after each step, are indicated by rectangles. The green shading represents a single processing step of collecting and collating 16S database sequences. (PDF 91KB)

**Supplementary File 29. Schematic representation of the pipeline used to filter publicly available *Streptomyces* genomes.** (PDF 59KB)

**Supplementary File 30. Sankey plot showing counts of taxonomic names in source databases, assigned at ranks from phylum to genus, to sequences identified with a key word ‘*Streptomyces*’ in the taxonomy field.** Note that Actinobacteria and Actinobacteriota are synonyms in LPSN for the correct Phylum name Actinomycetota, but that Actinomycetales and Streptomycetales are not taxonomic synonyms for each other. Streptomycetales is synonymous in LPSN with the correct name Kitasatosporales; Actinomycetales is a distinct taxonomic Order. The parent order of the Family Streptomycetaceae in LPSN is Kitasatosporales. (PDF 64KB)

**Supplementary File 31. Rectangular phylogram of the comprehensive maximum-likelihood tree of the genus Streptomyces based on the 16S sequence diversity of all 5,064 full-length 16S rRNA sequences with 100 TBE values.** (PDF 194KB)

**Supplementary file 32. Genomes sharing identical 16S rRNA sequences are assigned different names in NCBI. A total of 1,030 singleton clusters are not shown.** (PDF 8KB)

**Supplementary File 33. Phylogenetic tree.** PDF file showing collapsed phylogenetic tree showing distribution of members of the novel *Acintacidiphila* genus. (PDF 228KB)

**Supplementary File 34. Phylogenetic tree.** PDF file showing collapsed phylogenetic tree showing distribution of members of the novel *Phaeacidiphilus* genus. (PDF 228KB)

**Supplementary File 35. Phylogenetic tree.** PDF file showing collapsed phylogenetic tree showing distribution of members of the novel *Mangrovactinospora* genus. (PDF 228KB)

**Supplementary File 36. Phylogenetic tree.** PDF file showing collapsed phylogenetic tree showing distribution of members of the novel *Wenjunlia* genus. (PDF 228KB)

**Supplementary File 37. Phylogenetic tree.** PDF file showing collapsed phylogenetic tree showing distribution of members of the novel *Streptantibioticus* genus. (PDF 228KB)

## Introduction

Bacteria in the genus *Streptomyces* are among the most biotechnologically important groups of organisms and have been extensively screened for the production of bioactive metabolites[1]. *Streptomyces* are prolific producers of antimicrobial natural products, which changed the course of modern medicine in terms of treatment of infections and prophylaxis, and to prevent infections in immunosuppressed patients[2][3]. However, the use of these and other antimicrobials has led to the inevitable emergence of antimicrobial resistance (AMR) in the clinic[3]. AMR-related deaths are predicted to rise to 10 million annually and to become the leading cause of death worldwide in the by 2050[4]. The efficacy of current antibiotics is under threat, and discovery and development of novel medications is on the decline, so there is an urgent need to discover and develop novel antimicrobials for clinical applications [5]. Despite decades of investigation, recent genome mining efforts imply that a large number of natural products produced by the *Streptomyces* genus are yet to be identified and characterised, and it is hoped that among these may be found novel antimicrobials that could aid in combating the AMR crisis [2].

Exploring this anticipated range of potentially novel natural products has challenges. Empirical approaches have successfully delivered the majority of our pharmaceutically useful antimicrobials[6, 7], but they are labour intensive and often lead to rediscoveries of known antibiotics (the so-called dereplication problem)[8]. Further limitations arise from our inability to cultivate fastidious and recalcitrant microorganisms, and to synthesize bioactive compounds under typical laboratory conditions, which make empirical drug discovery less effective[7, 9]. In recent years a powerful strategy for drug discovery without the need for cultivating microorganisms has emerged involving mining of genome sequences for biosynthetic gene clusters (BGCs) predicted to be involved in the synthesis of natural products[10]. A number of bioinformatic tools for genome mining and bioprospecting, such as antiSMASH[3] GECCO[11], ARTS [12], PRISM[13], and numerous other tools based on a range of genetic and evolutionary conserved features, have been developed[14–16]. In conjunction with the commoditisation of genome sequencing, these tools have allowed the dereplication problem to be addressed rapidly and on a large scale to prioritise strains possessing potentially novel BGCs.

Despite the increased availability of whole genome sequences, 16S rRNA remains a standard approach for taxonomic identification, especially in the context of microbial communities, and metabarcoding. Clustering of 16S rRNA sequences into Operational Taxonomic Units (OTUs) at specified thresholds is a common approach for grouping similar sequences as a proxy for taxonomic identity[17]. Thresholds proposed in 1994, when only a small fraction of the currently-known 16S rRNA sequences were available, recommended that 95% identity be considered to bound representatives of the same genus, and 97% identity as a bound for species [17]. As bacterial genome sequencing became more common, it was realised that these thresholds were too permissive, and it has been common to treat each unique 16S rRNA sequence as a distinct taxonomic unit (zero-radius Operational Taxonomic Unit; zOTU)[18–20]. However, it is known that distinct species may share identical 16S rRNA sequences, including *Streptomyces* such as *S. ghanaensis* and *S. viridisporous*, and some species contain multiple copies of 16S rRNA with distinct sequences [21–23].

To maximise the potential of bioprospecting efforts through comparative genomics and pangenome analysis accurate information on the taxonomy of strains of interest is required[24]. *Streptomyces* taxonomy was historically based on polyphasic approaches incorporating morphological characteristics, phenotypic, and single-gene phylogenetic analyses[25]. These analyses often struggled to provide robust phylogenies within the group. Labeda et al.[23] demonstrated the sequence diversity of *Streptomyces* through a landmark 16S rRNA phylogeny, defining more than 130 lineages within the genus, though with some poor phylogenetic resolution due to the limited length and sequence variation of the 16S gene[23]. In particular, the 16S phylogeny proposed in [21] has been found to be inconsistent with subsequent whole-genome phylogenies and distance measures[26, 27]. For example, the 16S rRNA gene is unable to distinguish between members of the taxonomic Orders *Streptomycetales* and *Frankiales*, [28, 29]. To address these issues Labeda et al.[30] later augmented with their approach with Multi Locus Sequence Analysis (MLSA) to further resolve the phylogenetic relationships within the group [31].

The emergence of large-scale genome sequence data allows us to revisit phylogenetic and evolutionary relationships in complex taxa, and whole-genome distance measures play an increasingly important role in delineating species and genera. However, there remains no consensus on the interpretation of genus or species boundaries on the basis of whole genome similarity[32, 33]. A 95% ANI threshold is commonly used as an operational definition of species boundary, but interpretations of genome coverage can be difficult due to variation in genome size (affected for instance by genome expansion events, and presence or absence of plasmids) and comparisons between genomes of differing quality. Moreover, pairs of genomes with homology across less than 50% of their total genome length may even represent different candidate genera, regardless of ANI[34]. The emergence of genome-derived measurements such as digital DNA–DNA hybridization (dDDH), average nucleotide identity (ANI), average amino acid identity (AAI) and MinHash-related approaches has enabled rapid estimation of taxonomic relationships between organisms on the basis of their genomes [33, 35–38]. Utilising genomic data with these methods allows for incorporation of more genomic information, potentially offering higher phylogenetic resolution at the expense of averaging across multiple evolutionary processes [27, 39]. This has allowed the detection of mislabeled genomes, but it remains important to treat whole-genome taxonomy with some caution as mislabelled and low-quality (e.g. contaminated or incomplete) genomes may lead to erroneous results [40].

Here, we perform comprehensive classification of *Streptomyces* using 16S and whole-genome distance methods, to delineate the genomic landscape of this group of organisms, and to identify and interpret incongruences between the two approaches. We study all 48,981 available full-length 16S rRNA *Streptomyces* sequences and the 2,276 *Streptomyces* genomes available at the time of writing. Using 14,239 distinct full-length *Streptomyces* 16S rRNA sequences, we present the most comprehensive 16S phylogeny of *Streptomyces* known to us at the time of writing. We investigate the effect of clustering these 16S rRNA sequences into OTUs across a range of threshold identities including previously recommended 97% and zOTU identity thresholds. Finally, we examine the distribution of individual 16S rRNA sequences across *Streptomyces* genomes to determine whether a mapping exists between 16S rRNA sequences and whole genome-derived species boundaries, with a specific goal to understand whether zOTUs are congruent with genome-based classifications.

## Materials and Methods

### 1. Acquisition of 16S rRNA *Streptomyces* sequences from major 16S rRNA databases

16S rRNA sequences were manually downloaded from SILVA v138.1[41], RDP v11.5[42], NCBI[43] (New ribosomal RNA BLAST) PRJNA33175 and Greengenes v13.5[44] as indicated in *Table 1* (`Supplementary File 28`).

**Table 1.**
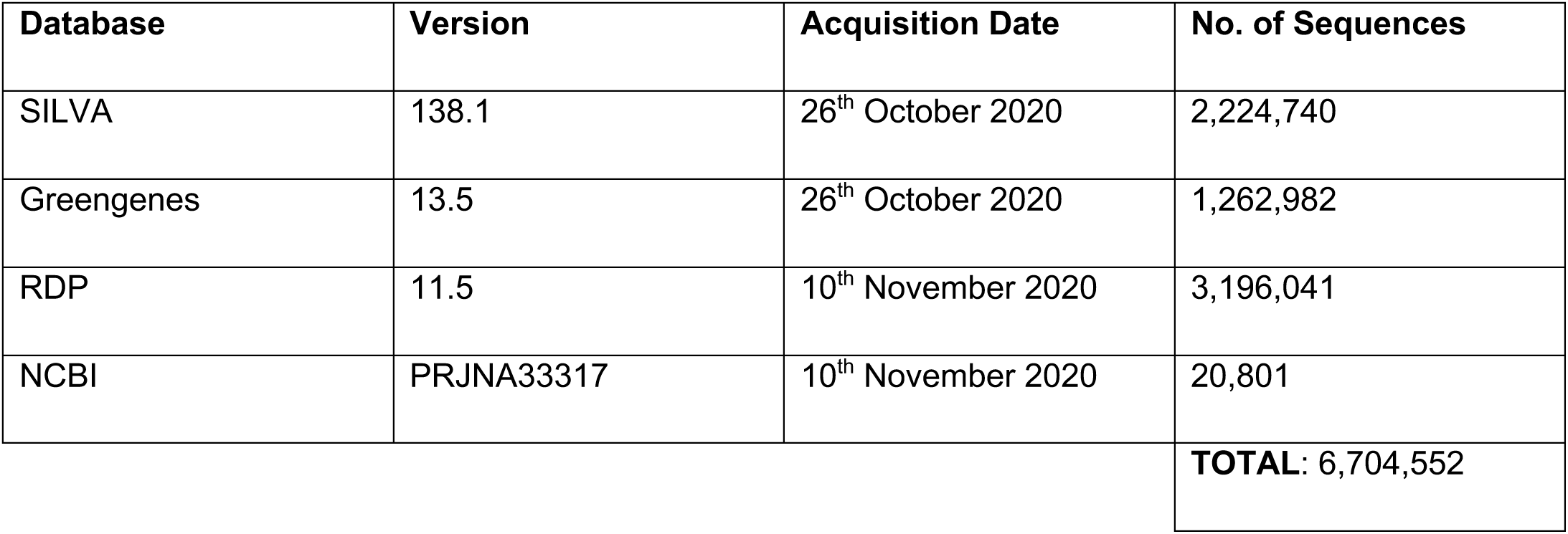
Summary description of 16S rRNA databases used in this manuscript.

| Database | Version | Acquisition Date | No. of Sequences |
| --- | --- | --- | --- |
| SILVA | 138.1 | 26<sup>th</sup> October 2020 | 2,224,740 |
| Greengenes | 13.5 | 26<sup>th</sup> October 2020 | 1,262,982 |
| RDP | 11.5 | 10<sup>th</sup> November 2020 | 3,196,041 |
| NCBI | PRJNA33317 | 10<sup>th</sup> November 2020 | 20,801 |
|  |  |  | **TOTAL: 6,704,552** |

Of these, Greengenes is the only database that stores sequence and taxonomy information in separate files. The taxonomy file (`gg_13_5_taxonomy.txt, Supplementary File 2`) was accessed on the 26th of October 2020, and taxonomic information was mapped to sequence by matching the sequence identifier (`gg_map_taxonomy.py; Supplementary File 2`).

### 2. Selection of full-length *Streptomyces* 16S rRNA sequences

To obtain full-length 16S sequence candidates, we filtered to retain only sequences with the keyword *Streptomyces* in the taxonomy field and excluding any 16S rRNA with length <1200bp. The resulting 48,981 sequences were combined to create a local 16S database (`all_DNA_whole_16S_strep.fasta, Supplementary File 3`), and base coding was standardised to thymine, replacing uracil (`get_complete_strep_seq.py, Supplementary File 3`).

### 3. LPSN Nomenclature Validation

Standardised nomenclature data was downloaded from the List of Prokaryotic Names with Standing in Nomenclature[45] (LPSN`; LPSN_taxonomy.csv, Supplementary File 6`) on 16^th^ February 2023. Species-level nomenclature previously assigned to the 48,981 full-length 16S rRNA *Streptomyces* sequences in the source database(s) was validated against this list (`get_NCBI_taxID_and_LPSN_status.py`; `Supplementary File 6).`

### 4. Elimination of nomenclature disagreements at higher taxonomic ranks

In some cases, the stated taxonomy at higher ranks of a sequence was a lineage not recognised to contain *Streptomyces spp.* The nomenclature at ranks up to Kingdom assigned to extracted full-length 16S rRNA *Streptomyces* sequences in the source database(s) were validated (`check_nomenclature_hierarchy.py`, `Supplementary File 3)` to identify and note, but not correct, this and similar cases of nomenclature hierarchy disagreement.

### 5. Removing redundant and ambiguous sequences

We removed (i) 24,132 redundant sequences identified using vsearch v2.15.1[46] (`remove_redundancy.sh`, `Supplementary File 4`), (ii) eleven sequences with more than 153 ambiguity symbols, (this threshold was determined by the negative binomial distribution of ambiguity symbols in the dataset - for detailed information see `remove_ambiguity` in `Supplementary File 4`), and (iii) 2,158 chimeric sequences identified using vsearch v2.15.1 (`remove_chimeras.sh`, `Supplementary File 4`). These operations reduced the local full-length 16S rRNA *Streptomyces* database (`all_DNA_whole_16S_strep.fasta`) to a total of 22,680 sequences (`non_chimeras.fasta, Supplementary File 4`).

### 6. Clustering of complete 16S rRNA *Streptomyces* sequences

The 22,680 retained full-length *Streptomyces* sequences were clustered using vsearch 2.15.1[46] at 30 evenly-spaced percentage identity thresholds ranging from 98% to 100% in steps of 0.1% (`cluster_sequences.sh, Supplementary File 5`). The number of distinct taxonomic assignments in a cluster for each clustering threshold was determined using the NCBI reference taxonomy[47] (downloaded on the 31^st^ January 2023 from https://ftp.ncbi.nih.gov/pub/taxonomy/taxdmp.zip; `names.dmp, Supplementary File 6`). Each sequence was assigned NCBI taxID corresponding to the LPSN-validated nomenclature assigned to it in the source database(s). Nomenclature and corresponding taxIDs for redundant sequences removed in Methodology Section 5 were included in the analysis (`cluster_composition_analysis.py, Supplementary File 6`).

### 7. Phylogenetic reconstruction

All unique 16S rRNA sequences (zOTUs), and a further ten 16S rRNA outgroup sequences belonging to isolates from *Kitasatsopora, Streptoalloteichus* and *Clavibacter* genera (`outgroup.fasta, Supplementary File 9`) were aligned using nextalign v0.1.4[48] against the GCF_008931305.1 16S rRNA reference sequence (`S_coelicolor_A32.fasta, Supplementary File 9`) from *S. coelicolor* A3(2). Alignments were trimmed using trimAl v1.4[49] (`trim_alignments.sh, Supplementary File 9`), and subsequently dereplicated using RaxML (`alignment_dereplication.sh, Supplementary File 9`). A Maximum-Likelihood tree was estimated using RaxML-NG v1.0.2[50] with 100 Transfer Bootstrap Expectation (TBE) replicates (`raxml_bootstraps.sh; raxml_tbe.sh, Supplementary File 10`) on the ARCHIE-West computing cluster with Intel XI(R) Silver 4216 CPU 2.10Hz, 32 cores and 187 GB RAM.

### 8. Assessment of unique 16S rRNA sequences from *Streptomyces* genomes

All 2,276 publicly available *Streptomyces* genome sequences (`streptomyces_genomes.txt, Supplementary File 17`) were downloaded from NCBI on July 8^th^, 2021 (`download_genomes.sh`, `Supplementary File 17`). Their assembly status was checked against the NCBI assembly report downloaded on 30^th^ January 2023 from https://ftp.ncbi.nlm.nih.gov/genomes/refseq/assembly_summary_refseq_historical.txt (`assembly_summary_refseq_historical.txt, check_genome_status.py, Supplementary File 17`). We discarded 120 suppressed genomes from the analysis. Updated versions of five replaced genomes (`streptomyces_replaced_genomes.txt, Supplementary File 17`) were manually downloaded from NCBI[51] on 30^th^ January 2023. A total of 6,692 16S rRNA sequences were extracted from the 2,156 *Streptomyces* genomes by matching the key word ‘16S’ in the `\gene_qualifiers` GenBank field (`extract_16S.py, Supplementary File 18`). We filtered the extracted sequences to a total of 4,227 by retaining only the 1,369 genomes (`retained_genomes.txt, Supplementary File 18`) that exclusively contain only full-length and ambiguity symbol-free 16S rRNA sequences (`filter_16S_seq.py`, `filtered_16S_seq_from_strep_genomes.fasta, Supplementary File 18`). All 4,227 such sequences extracted from *Streptomyces* genomes were aligned using nextalign v0.1.4[48] against the same GCF_008931305.1 16S rRNA reference sequence as used previously (`S_coelicolor_A32.fasta, Supplementary File 9`). The alignment was trimmed using trimAl v1.4[49] (`trim_alignment.sh, Supplementary File 19`), and genomes sharing identical 16S rRNA sequences were clustered (`get_input_genomes_for_pyani.py, Supplementary File 19`) and used as input sequences to determine their taxonomic boundaries with ANI using pyANI v0.3[34] (`pyani_analysis.sh, Supplementary File 19`). We adopted the 50% coverage and 95% identity thresholds to estimate genus and species boundaries, respectively. ANIm analysis using pyANI v0.3[34] was performed to determine taxonomic boundaries for all genomes found in the same connected components (`pyani_analysis.sh`, `Supplementary File 20`).

### 9. Network analysis of genomes based on shared 16S rRNA sequences

To visually represent connections between the 1,369 *Streptomyces* genomes that contain only full-length and ambiguity symbol-free 16S rRNA sequences, we constructed a network representing individual genomes as nodes, and assigning edges between genomes with weights corresponding to the number of shared identical 16S sequences, the graph being processed using NetworkX[52], and visualised with plotly v5.6.0 (https://plotly.com/python/; `genome_16S_NetworkX.ipynb; Supplementary File 20`). Edges corresponding to pairs of genomes with no 16S sequence in common were removed prior to construction of the graph. Node layout was calculated using Cytoscape[53] v3.9.0 with Prefuse Forced Directed layout.

## Results and Discussion

### Public 16S *Streptomyces* sequence databases include records with low-quality or redundant sequence, or that have issues with taxonomic nomenclature

High-quality sequences are crucial for ensuring the accuracy and reliability of research and analysis, particularly in the fields of genomics, taxonomic classification, and applied microbiology. Using unintentionally inaccurate database records can potentially lead to false interpretations and flawed research outcomes. Of the 62,482 16S rRNA sequences belonging to the genus *Streptomyces* across the SILVA v138.1, RDP v11.5, Greengenes v13.5 and NCBI (PRJNA33175) databases (Methodology Section 2), we determined that only 48,981 (78.4%) were full-length sequences. In total, we identified 24,849 non-redundant full-length 16S rRNA sequences (50.7% of all full-length *Streptomyces* sequences; 39.8% of all database *Streptomyces* sequences `Supplementary File 28`).

Prokaryotic taxonomic nomenclature is a pivotal mechanism for unambiguous communication about an organism’s identity. Correct nomenclature helps avoid undesirable clinical, ecological, agricultural, and pharmaceutical consequences[54, 55]. The databases considered in this work rely on a variety of taxonomic authorities: RDP uses Bergey’s Manual[56, 57], SILVA uses LPSN and Bergey’s Manual[45]; and NCBI combines nomenclature provided by the submitter with that in Greengenes, basing its nomenclature on that in the NCBI taxonomy[44, 58]. All of these schemes are good-faith efforts to follow the International Code of Nomenclature of Prokaryotes (ICNP)[59], but we find that records are in some cases inaccurate. LPSN is an online database that catalogs the validly published names of prokaryotes in accordance with the Rules of ICNP[45]. We validated taxonomic nomenclature assigned to each of the full-length sequences in the source database(s) against LPSN (Methodology Section 3) and find that only 14,859 (30.3%) of the 48,981 species names assigned to extracted full-length *Streptomyces* sequences were found within LPSN. A further 1,400 (2.9%) were synonyms of valid names, but 28,333 (57.8%) sequences were labelled as unclassified *Streptomyces*, 17 (0.03%) were misspelled, and no record in LSPN was found for 4,372 (8.9%) sequences. The LSPN status of all 48,981 sequences used in this manuscript is provided in `full_length_strep_records_info.csv` (`Supplementary File 6`).

In the originating databases, sequences may be annotated with synonyms of taxon names at various ranks, and it is reasonable to expect that these taxon names should be consistent within the same lineage. Our investigation of nomenclature within the source databases at ranks from Kingdom to Genus (Methodology Section 4) identifies sequences having the *Streptomyces* keyword in the taxonomy field that are assigned: (i) correctly to all ranks above *Streptomyces*, eg. AWQW01000120.100.1367 belongs to *Bacteria* kingdom, *Actinobacteriota* phylum, *Actinobacteria* class, *Streptomycetales* order, *Streptomycetaceae* family, *Streptomyces* genus and *Streptomyces niveus* species (we note that nomenclature is fluid and, for example, at phylum level *Actinobacteriota* has been superseded by *Actinomycetota* [60], but choose to reflect the assigned nomenclature in the database); (ii) to higher ranks not expected to contain the *Streptomyces* genus (eg. KY753270.1.1450 belongs to *Bacteria* kingdom, *Firmicutes* phylum, *Bacilli* class, *Bacillales* order, *Bacillaceae* family, *Bacillus* genus, and *Streptomyces pseudovenezuelae* species); (iii) to higher ranks that correctly include *Streptomyces* from kingdom to genus, but where the annotated nomenclature nevertheless disagrees on the parent genus name (eg. JN987181.1.1444 belongs to *Bacteria* kingdom, *Actinobacteriota* phylum, *Actinobacteria* class, *Streptomycetales* order, *Streptomycetaceae* family, *Streptomyces* genus and *Lactobacillus apodemi* species); and (iv) to ambiguous hierarchies where there is a complete lack of information about higher ranks (eg. NR_042095.1). This is consistent with previous observations of taxon names assigned to conflicting nomenclature at higher ranks in Greengenes and SILVA[61]. All identified nomenclature at ranks from kingdom to genus are summarised as a Sankey plot in `Supplementary File 30`, and a complete list of all 48,981 investigated sequences is given in `nomenclature_hierarchy_info.csv` (`Supplementary File 3`).

Chimeric sequences, and sequences with a high proportion of ambiguity symbols, can be destructive in phylogenetic analyses, leading to model misspecification, incorrect branch length and topology estimation[62]. Eleven sequences with more than 153 nucleotide ambiguity symbols were discarded from our dataset prior to analysis. During the data cleaning process, a further 2,158 potential chimeric sequences were also identified and removed from the dataset. Following the filtration and cleaning process, 22,680 full-length non-redundant high-quality sequences (46% of the initial dataset) were taken forward for further analyses and phylogenetic tree reconstruction. Despite significant and diligent long-term efforts by curators to remove poor quality sequences from the databases used in this manuscript[44], we still required to identify a large amount of data to avoid introducing identifiable sources of potential inaccuracy to our analysis (Methodology Section 5).

### A comprehensive *Streptomyces* 16S phylogeny

To estimate the evolutionary relationship amongst *Streptomyces* all 14,239 zOTU sequences were used to produce a multiple sequence alignment (MSA; Methodology Section 8). Ten outgroup sequences from related non-*Streptomyces* genera were added to the analysis to aid in root placement (`Supplementary File 9`). The MSA was trimmed to 1,086 nucleotides and 9,049 sequences after redundant sequences were removed (no positions in the alignment were absolutely conserved), and a maximum-likelihood tree was calculated (Methodology Section 7). Clades containing a single species taxonomic assignment were collapsed to single leaf nodes to facilitate visualisation of the phylogenetic tree for all 5,064 nodes shown in **Fig. 1** (full tree provided in newick format in `Supplementary File 10; 04_TBE.raxml.support`; collapsed newick file version provided in `Supplementary File 11; collapsed_strep_tbe.new`).

**Fig. 1.**
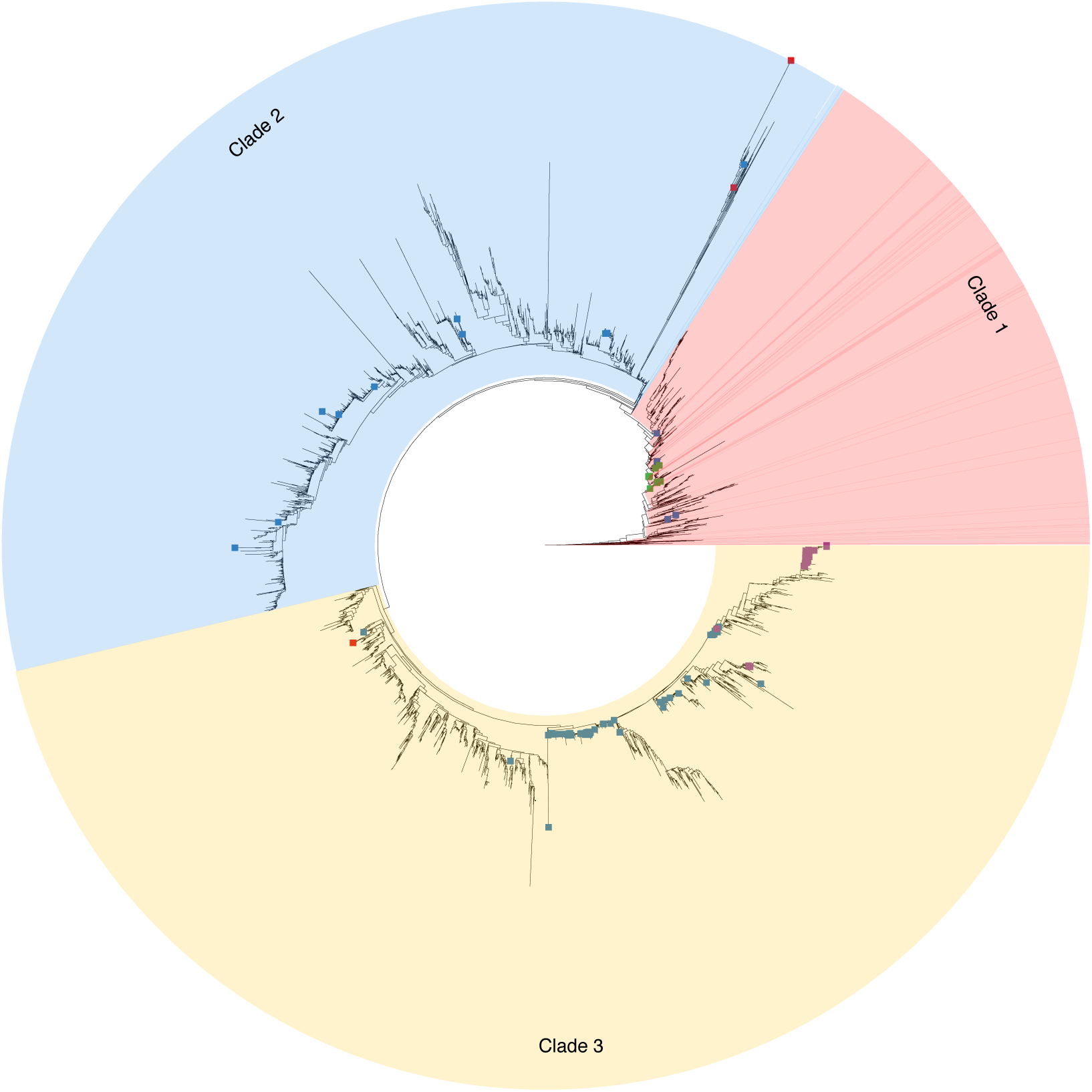
Maximum-likelihood tree of the genus *Streptomyces* constructed from 9,049 full-length 16S rRNA sequences. Clades containing a single assigned species-level taxon were collapsed to single leaf nodes. Three major clades are highlighted in distinct colours. Squares indicate distributions of 16S sequences assigned the same species names in the source database(s); *S. griseus* sequences are shown in blue, *S. clavuligerus* in red, *S. lydicus* in green, and *S. scabei* in purple. Sequences with these species assignments tend not to be monophyletic, indicating incongruence between taxonomy and the 16S gene tree. An equivalent rectangular phylogram is provided in `Supplementary File 31`.

To our knowledge this is the largest, most comprehensive 16S rRNA phylogenetic reconstruction attempted for *Streptomyces* to date. No clade received TBE support greater than 60%, and only 18 clades have a TBE support above 50%; hence we do not consider the topology of this tree to be robust as presented. `Supplementary File 12` describes a phylogenetic tree where clades having TBE value of 50% or higher are marked.

A notable feature of the ML tree is the presence of three clearly distinguishable major clades (**Fig. 1**). If 16S sequence databases were simply consistent with the true *Streptomyces* species tree, we might expect to find that 16S sequences taxonomically assigned to the same species are broadly monophyletic (allowing for ancestral duplications, given that a large proportion of *Streptomyces* species contain multiple 16S loci). If the 16S tree departs from this distribution of taxonomic assignment, it might reflect an unreliability in classification of taxa using the existing database assignments. To examine this, we mapped pharmaceutically and agriculturally important isolates to the comprehensive 16S ML phylogeny in figure 1. We find that some species annotations are consistently found within a single clade (*S. lydicus, S. albulus* and *S. venezuelae*, `Supplementary File 14`), and some exhibit dispersion within major clades consistent with limited misannotation (*S. scabei*, *S. lavendulae* and *S. rimosus* (`Supplementary File 16`). However, we find some taxon representatives to be distributed widely across the tree, in two or more major clades (*S. griseus, S. albus:* `Supplementary File 13`). Other species assignments are represented by an insufficient number of sequences to clearly demonstrate a pattern (*S. clavuligerus* or *S. coelicolor:* `Supplementary File 13`). Additionally, recent reclassifications within *Streptomycetaceae* family lead to the proposal of six novel genera[31]. To determine whether the proposed novel genera resolve in our analysis, we checked their distribution across the 16S ML phylogeny. We find that members of *Wenjulia* (`supplementary file 36`) are consistently placed on the comprehensive 16S ML phylogeny, whereas members of *Actinacidiphila* are scattered across the tree (`supplementary file 33`). Additionally, our analysis did not include any representative from *Peterkaempfera* genus, and the available sequences for *Mangrovactinospora* (`supplementary file 35`), *Phaeacidiphilus* (`supplementary file 34`) and *Streptantibioticus* (`supplementary file 37`) were insufficient to establish a clear pattern in their distribution. Overall we find evidence of taxonomic misassignment across the full scope of 16S sequences, consistent with observations of sequence mis-annotation previously estimated for SILVA and GreenGenes to be around 17% at ranks up to phylum, and a similar mis-annotation rate of 10% in RDP[61]. It is possible that, in some cases, the apparent dispersion of a single taxon across the tree could the result of limited sequence variation within the 16S rRNA and failure to obtain a robust phylogeny, given that most internal nodes have a TBE support value of lower than 50% (**Fig. 1**). However, the three major clades do appear to be relatively robustly distinguished in the phylogeny, and where the same taxon has representatives in two or more of these clades we consider that this calls into question the corresponding assignment.

### 16S percentage sequence identity thresholds do not reliably delineate existing *Streptomyces* species assignments

The long-standing 16S rRNA clustering threshold for species separation of 97% sequence identity has been robustly questioned[63], and current best practice is to use zOTUs, or Amplicon Sequence Variants (ASVs) for taxonomic classification and clustering of 16S and other marker genes. To identify whether any clustering threshold adequately circumscribes *Streptomyces* taxa, we applied a range of threshold identities to over 22,000 full-length *Streptomyces* 16S rRNA sequences, then determined the agreement between taxonomic species and cluster membership (Methodology Section 6). If zOTUs were assumed to be an accurate proxy for species, then with 14,239 known zOTUs one might conclude that there are the same number of candidate *Streptomyces* species. Currently at least 650 species are recognised within *Streptomyces*, and so many observed zOTUs might suggest either a high degree of cryptic species diversity, or that 20 or more distinct 16S rRNA sequences may be characteristic of the same *Streptomyces* species. Our initial examination of pharmaceutically important strains *S. griseus* (streptomycin producer), *S. lydicus* (natamycin, lydimycin and streptolydigin producer), *S. clavuligerus* (clavulanic acid and cephamycin C producer[1]) and the phytopathogen *Streptomyces scabiei*[64] shows that 16S rRNA sequences assigned these species names are split across multiple zOTUs. Specifically, *S. clavuligeus* is split across 8 zOTUs, *S. griseus* is found in 145 zOTUs, *S. lydicus* in 16, and *S. scabiei* in 62. These findings are consistent with observations *Streptomyces* genomes possess multiple non-identical 16S rRNA sequences (`Supplementary File 26`). The existence of multiple distinct 16S rRNA sequences corresponding to a single species implies that naïve 16S metabarcoding of communities assuming that zOTU diversity reflects species diversity may overestimate the number of species per sample.

As we raise the 16S sequence identity clustering threshold from 98% to 100%, clusters increase in number and tend to have fewer 16S sequence members (`Supplementary File 7`). The number of unique taxa per cluster also tends to fall as the identity threshold approaches 100% **(Fig. 2)**. We identified 10,548 zOTUs containing two or more sequences. Of these, 8,326 (78.9%) included at least one sequence currently named as *Streptomyces* sp. whose classification is ambiguous, and 4,820 (45.7% of zOTUs) consisted only of such unclassified sequences. This significant proportion of unclassified sequences might correspond to distinct species or represent as yet unclassified members of known species, including as other species represented in the cluster. These sequences introduce ambiguity and disrupt estimation of the true taxonomic accuracy of 16S marker sequences.

**Fig. 2.**
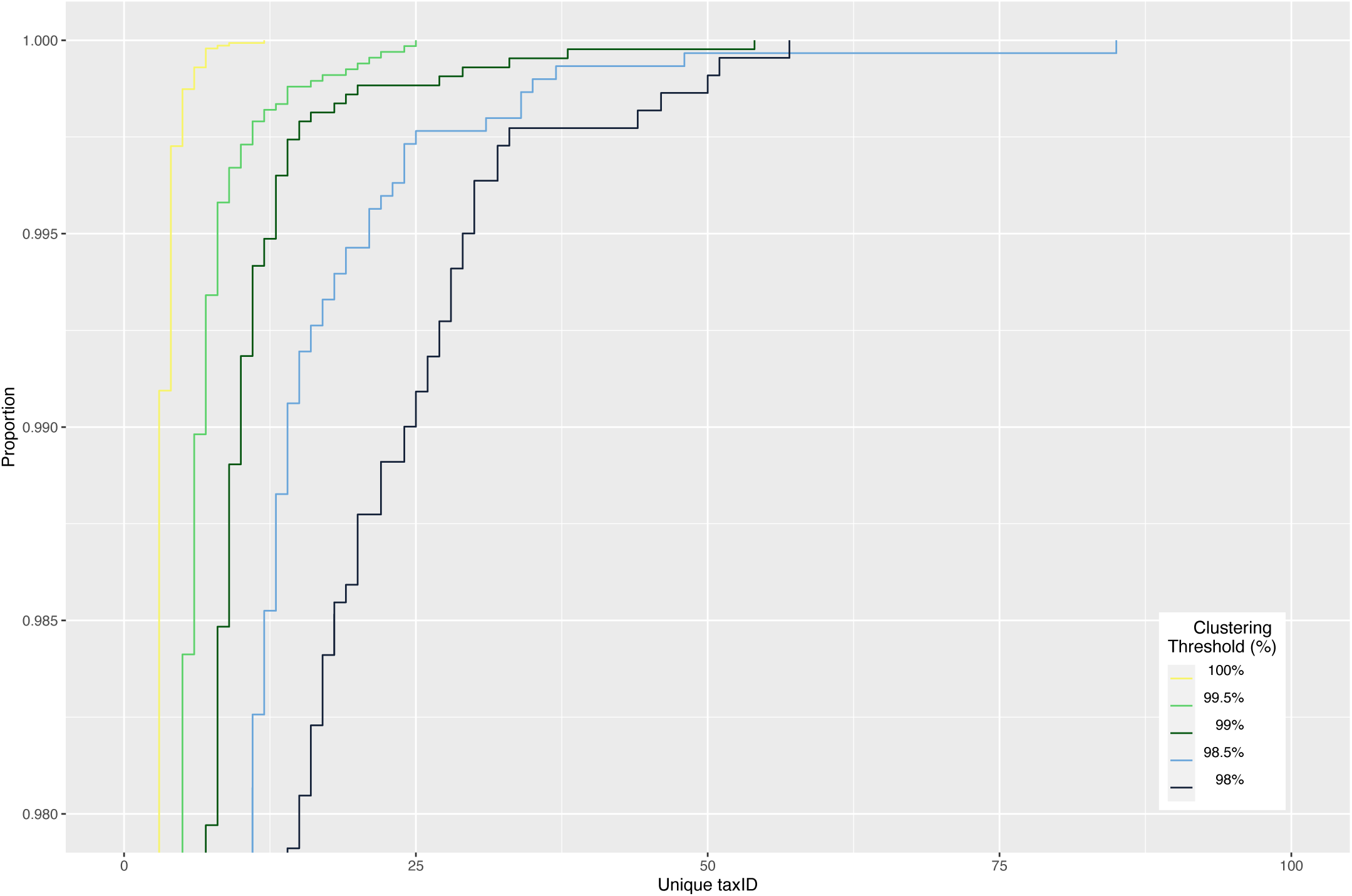
Empirical Cumulative Distribution of taxonomic composition at varying %ID thresholds.

Nevertheless, if 16S rRNA sequences do provide sufficient sequence diversity to distinguish between *Streptomyces* species, and zOTUs are a reliable proxy for taxonomic assignment at or below species-level in *Streptomyces,* then each zOTU cluster should contain only a single unambiguous species name. However, our analysis shows that 3,747 (26%) of zOTUs contain 16S sequences currently assigned to at least two distinct *Streptomyces* species (`Supplementary File 8;cluster_taxID_info.csv, Supplementary File 6`). One zOTU notably includes sequences annotated as twelve distinct species (*S. coelicolor, S. albidoflavus, S. somaliensis, S. rutgersensis, S. paulus, S. limosus, S. griseochromogenes, S. sampsonii, S. resistommycificus, S. felleus, S. violascens* and *S. sp.*). We therefore find that a substantial fraction of full-length 16S zOTUs do not map exactly to a single *Streptomyces* species assignments, and in particular single 16S sequences frequently map to multiple distinct species.

### Whole-genome sequence classification indicates that distinct Streptomyces species can share identical full-length 16S sequences

No strain information linking to sequenced genomes could be recovered for 16S rRNA sequences from SILVA, Greengenes, RDP or NCBI new ribosomal RNA BLAST databases. Therefore, to map the 16S sequence landscape to the whole-genome classification landscape, we extracted 6,692 16S rRNA full-length sequences from 2,156 publicly available *Streptomyces* genomes (Methodology Section 9). Eighty-seven of the published assemblies lacked identifiable 16S rRNA sequences, and 700 genomes contained at least one 16S rRNA sequences with ambiguity symbols or partial sequences that could lead to biased observations and overestimation of the intragenomic diversity of 16S rRNA sequences. These genomes were excluded from our analysis, yielding a dataset comprising 4,227 16S rRNA sequences from 1,369 genomes containing only full-length, ambiguity symbol-free 16S rRNA sequences.

*Streptomyces* genomes most commonly contain six copies of 16S rRNA operons[65]. Across the 1,369 assemblies analysed we find that 16S rRNA sequence copy number varies from one to twelve copies per genome (`Supplementary File 26`). It was found that 359 (26.2%) of the assemblies contained six copies, 144 (10.5%) had more than six copies, and 865 (63.1%) contained fewer than six copies of 16S rRNA. The genomes of 375 (27.4%) assemblies were found to contain multiple non-identical 16S rRNA sequences. A single assembly (GCF_900199205.1) was found to contain eight distinct 16S rRNA sequences but the majority, 993 genomes (72.5%), possessed only a single 16S rRNA sequence variant, of which 811 contained only one detectable copy of 16S rRNA. Inconsistency in the number of 16S rRNA operons and their intragenomic heterogeneity could be due to a range of causes, but is likely due to inclusion of *Streptomyces* genomes assembled to different levels of completeness and quality (eg. contig, scaffold, complete and chromosome). Assemblies could also have been affected by the presence of duplication artifacts or collapsed 16S rRNA sequences resulting from sequencing errors, such as overassembly of short reads from distinct 16S rRNA loci into a single artifactual 16S sequence. It may be assumed that NCBI complete and chromosomal *Streptomyces* assemblies reflect the true genomic heterogeneity of 16S rRNA sequences. Given this, it would be expected that 69% of *Streptomyces* isolates would contain multiple distinct 16S rRNA sequences (`Supplementary File 27`). This proportion is consistent with previous observations that *Streptomyces* strains may contain multiple distinct 16S copies, but twice that observed across all *Streptomyces* genome assemblies in NCBI [66].

To further investigate the potential for inaccurate *Streptomyces* taxonomic assignment when using 16S rRNA sequences, we examined the relationship between distinct genomic 16S sequences and the species assignments of their corresponding genomes. We examined the distribution of 16S sequences by the number of genomes they occur in, and the number of uniquely assigned species names in NCBI associated with those genomes (`Supplementary File 32`). We find that a single 16S sequence variant may be represented in as many as 33 genomes, and be associated with a group of genomes assigned as many as six species names.

Previous whole-genome analyses of *Streptomyces* also observed that identical 16S rRNA sequences are present in strains assigned to different species[67]. Some of the discrepancy may arise from differing approaches to, and knowledge of, taxonomic assignment over time that leads to, for example, the same strain being assigned to a different species depending on when the analysis was done. However, some of these observations may genuinely reflect common 16S sequence shared across sequence boundaries, as 16S substitution rates may be slow in relation to speciation events.

For *Streptomyces* species multiple distinct 16S rRNA sequences can be found within the same genome (`Supplementary File 26`), implying that there is a one-to-many mapping between *Streptomyces* species and 16S rRNA sequence. It follows that it is not always possible to cluster *Streptomyces* 16S sequence data without splitting a single organism into multiple zOTUs. Simple counts of *Streptomyces* zOTUs when metabarcoding with 16S may thus overestimate species numbers. It also follows that comprehensive 16S rRNA gene trees reflect gene histories, and may not directly recapitulate the corresponding accurate species trees.

We constructed network graphs from genome-derived 16S sequences to visually represent connections between *Streptomyces* genomes based on their common 16S sequences, and thereby interpret the relationship of this network to whole-genome similarity based species classifications (Methodology Section 9). We represented each of 1,369 Streptomyces genomes as a node in the graph, connecting two genomes with an edge if they shared an identical full-length 16S rRNA sequence. Our network analysis resolved the 1,369 *Streptomyces* genomes into 709 connected components (**Fig. 3**). The largest connected component united 47 genomes, but 527 (74.3%) genomes were singletons, sharing no 16S sequence with any other *Streptomyces* assembly (an interactive version of this graph is provided in `Supplementary File 22`).

**Fig. 3.**
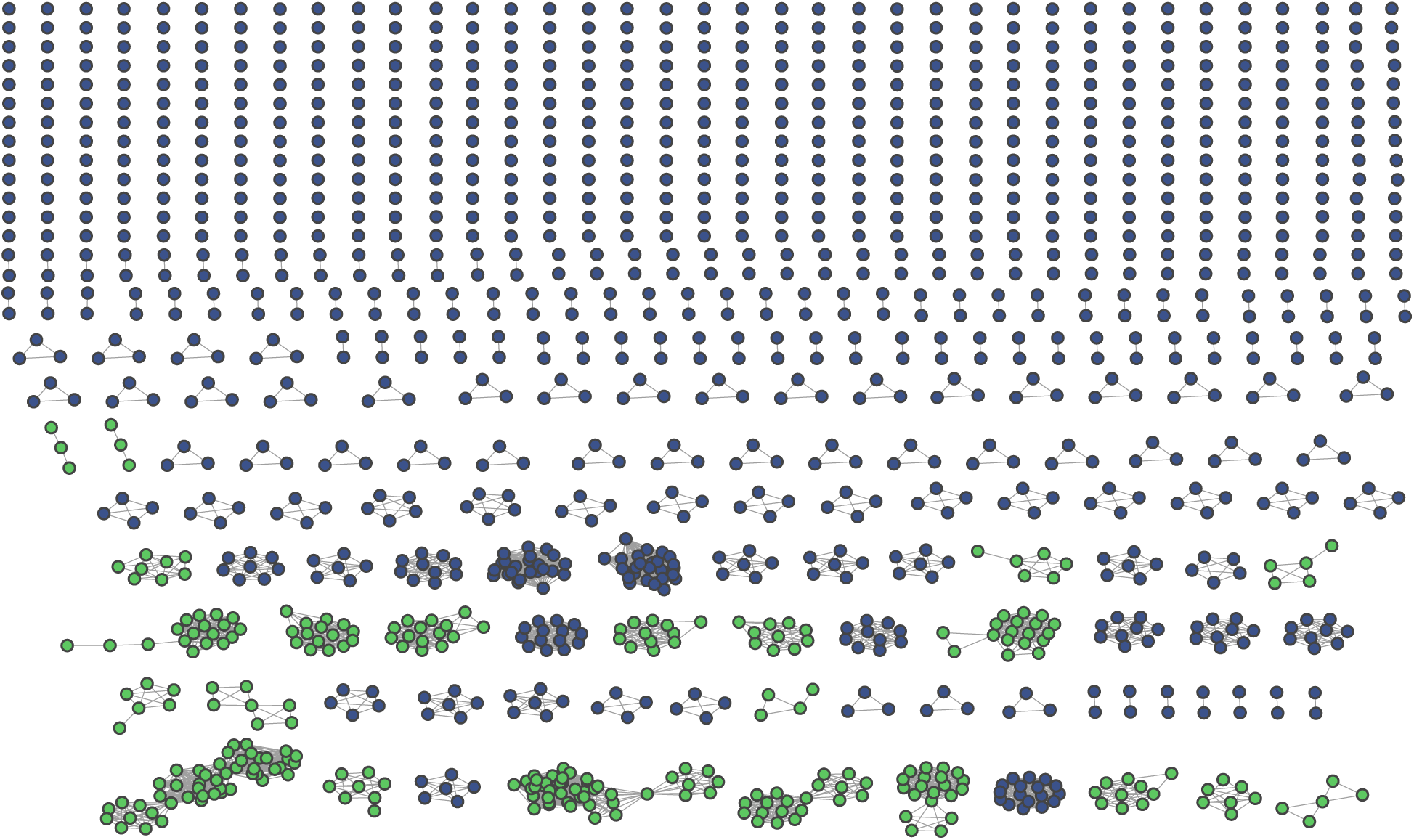
Network graph with 1369 genomes and 709 connected components. Each node represents a distinct genome assembly, each edge corresponds to at least one identical 16S sequence being shared between that pair of genomes. Blue connected components form cliques, in which every genome shares at least one identical 16S sequence with all other genomes in the same connected component. Green connected components do not have this property.

If there was a direct mapping between 16S rRNA sequences and species, then connected components would be expected to form cliques (*k*-complete graphs) where each genome within a single connected component would be linked to every other genome in the same component by at least one edge. However, 22 connected components formed non-cliques (**Fig. 3**), indicating that some genomes within a single connected component may not share any identical 16S sequence with some of the other genomes in the same component. If all members of a component belong to the same taxon, this would imply that two members of the same taxon might share no 16S sequence in common. Alternatively, some genomes from distinct taxa may share identical 16S rRNA sequences, perhaps resulting in multiple species being found within a single connected group of genomes. Such a situation might arise for a number of reasons, including inter-species recombination[68, 69] or selective pressures from natural products that act upon the ribosome[70].

We performed ANI analysis on the genomes comprising each connected component (Methodology Section 9) to establish whether the subgraph corresponded to a single grouping of genomes at genus or species level. We defined genomes as belonging to the same candidate genus if they shared at least 50% genome coverage, and belonging to the same species if they shared at least 95% ANI. These boundaries are approximations, but correspond to commonly-used heuristics[33]. We did not find that connected components always represent only a single kind of taxonomic relationship between its members. Instead we observe components comprising: (i) a single species, where all genomes share at least 50% genome coverage (**Fig. 4A**) and 95% ANI (**Fig. 4B**); (ii) a single genus, but multiple species, where all genomes share at least 50% genome coverage (**Fig. 4C**), but not all share at least 95% ANI (**Fig. 4D)**; (iii) potentially multiple genera, where some genomes share less than 50% genome coverage (**Fig. 4E**) and less than 95% genome identity (**Fig. 4F**). ANIm coverage and identity plots for the remaining connected components are provided in `pyani_heatmaps` folder (`Supplementary File 20`), and the numbers of subgraphs falling into each category are summarized in **Table 2**.

**Fig. 4.**
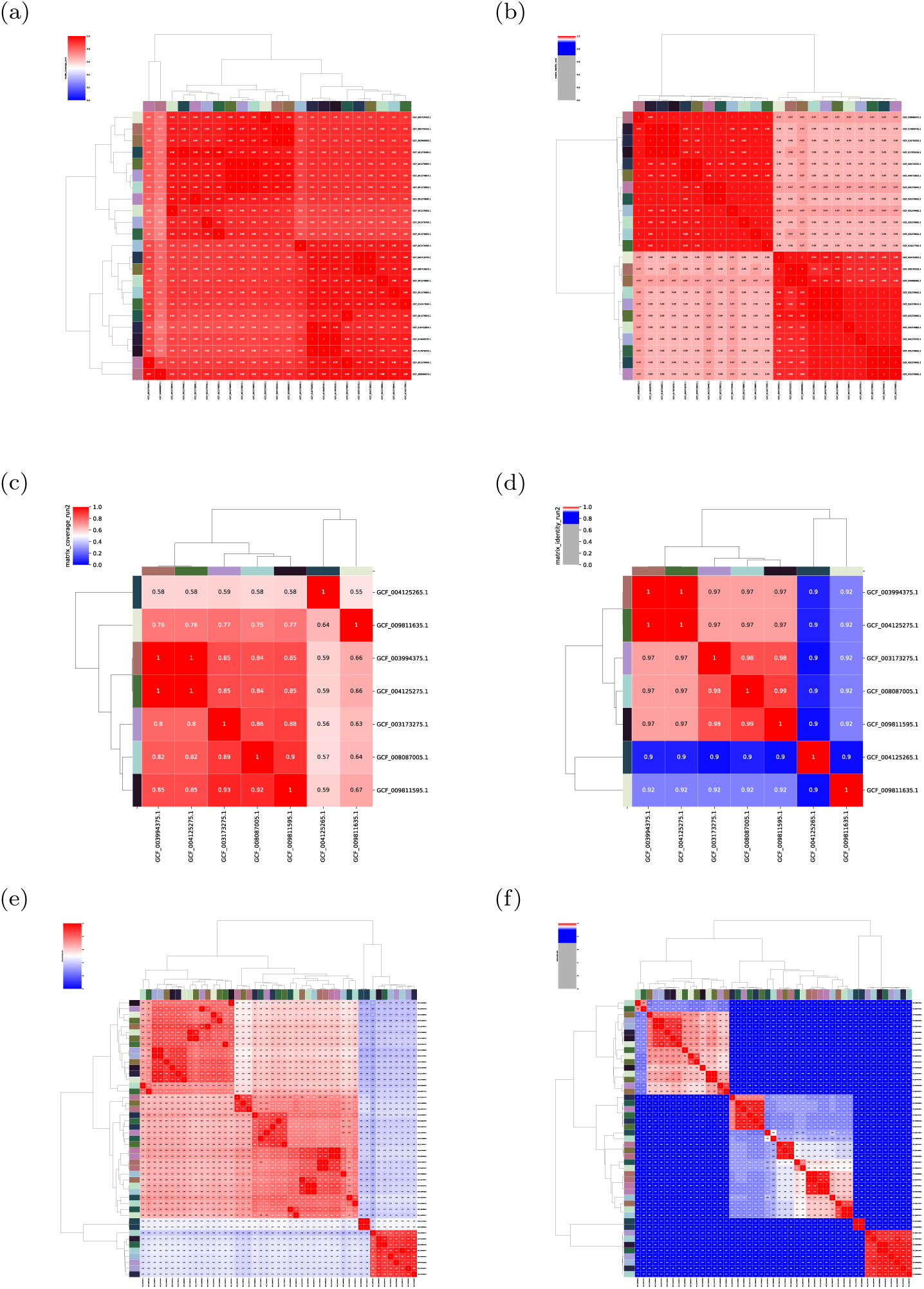
Heatmaps of ANIm coverage (left), and ANIm identity (right) for three example connected components. Heatmaps in the same row correspond to comparisons for the same connected component. The left column represents percentage genome coverage, the right column %ANI. Red cells in coverage plots correspond to genome coverage of 50% or above, interpreted as common membership of the same genus; blue cells correspond to coverage below 50% and imply distinct genus assignments. In ANI plots, red cells correspond to genome identity of 95% or above, interpreted as membership of the same species; blue cells represent imply distinct species. In some cases ANIm species classifications map onto components containing genomes from a single genus and species (a-b), to distinct species in the same genus (c-d), or to distinct genera (e-f).

**Table 2.** Summary statistics for taxonomic composition of subgraphs uniting at least two genome assemblies.

| Category | Count | Percentage |
| --- | --- | --- |
| >≈50% coverage; >≈95% identity | 154 | 84.62% |
| >≈50% coverage; <95% identity | 25 | 13.74% |
| <50% coverage; <50% identity | 3 | 1.65% |

By mapping ANIm coverage to visually represent the distribution of unique candidate genera within each connected components, we find that 174 (98.35%) non-singleton subgraphs unite isolates that share at least 50% genome coverage, indicating likely membership of the same genus (`Supplementary File 23`). However, three connected components (**Fig. 5A**) appear to comprise assemblies from distinct candidate genera. Using our whole-genome comparison threshold to define genus, and although the prevalence of such groups is low, we find that full-length 16S sequences are not always sufficient to resolve *Streptomyces* at genus level. We find, with a similar analysis using %ANIm identity (`Supplementary File 24`), that the majority (84%) of connected components likely represent a single species. However, 28 connected components (**Fig. 5B**) contain assemblies representing multiple species.

**Fig. 5.**
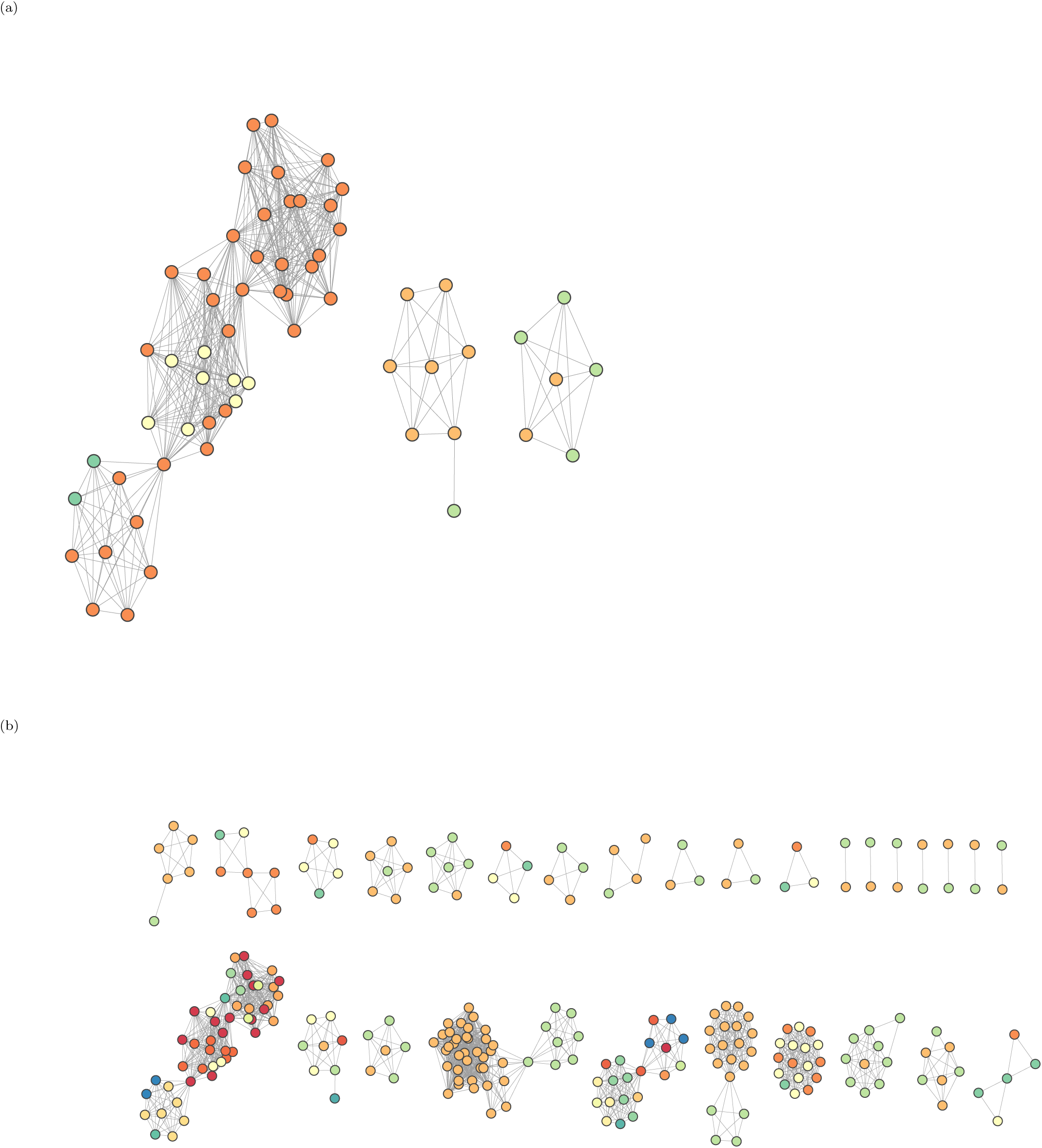
Connected components connecting assemblies from distinct candidate genus (A) and species (B). Nodes represent distinct genomes assemblies, while edges indicate the presence of at least one identical 16S sequence shared between the corresponding pair of genomes. Each unique candidate genus (A) or species (B) is represented as a distinct node colour.

Our data are consistent with previous observations made by Chevrette el al., [27], that distinct *Streptomyces* species confirmed by ANI may share identical 16S rRNA sequences. Moreover, our results also indicate that some networks of genomes which would be assigned as the same species using ANI do not form cliques linked by identical 16S sequences, and so it is possible that genomes assigned to the same *Streptomyces* species by whole-genome methods may not share *any* identical 16S sequences. Thus, in *Streptomyces*, there is a one-to-many mapping from Streptomyces species to 16S sequence, and a one-to-many mapping from 16S sequence to species. Taken together, our results demonstrate that use of 16S rRNA sequences in isolation for taxonomic classification of *Streptomyces* (as is often the case in 16S metabarcoding) can lead to a significant minority of misclassifications not just at the species level, as might be expected, but also at genus level.

To further delineate the relationship between 16S sequence variation and whole genome taxonomy, we measured relatedness within each 16S zOTU using ANIm (Methodology Section 9). We define a single zOTU genome cluster as a group of at least two genomes sharing an identical, full length and ambiguity symbol-free 16S rRNA sequence (`Supplementary File 32`). We classified pairs of genomes as belonging to the same genus if they share at least 50% ANIm coverage, and belonging to the same species if they share at least 95% ANIm identity, as before. We show the pairwise comparison results as 1D scatter plots of pairwise genome coverage (**Fig. 6**) and pairwise genome identity for each zOTU (**Fig. 7**). In the figure we subdivide zOTUs into groups corresponding to the number of distinct species currently assigned to the *Streptomyces* genomes containing that OTU (zOTUs used in this analysis contain between one and six distinct assigned *Streptomyces* species). We further overlay whole-genome comparison information by colouring comparisons differently if the participants correspond to distinct genera or species by our ANIm thresholds.

**Fig. 6.**
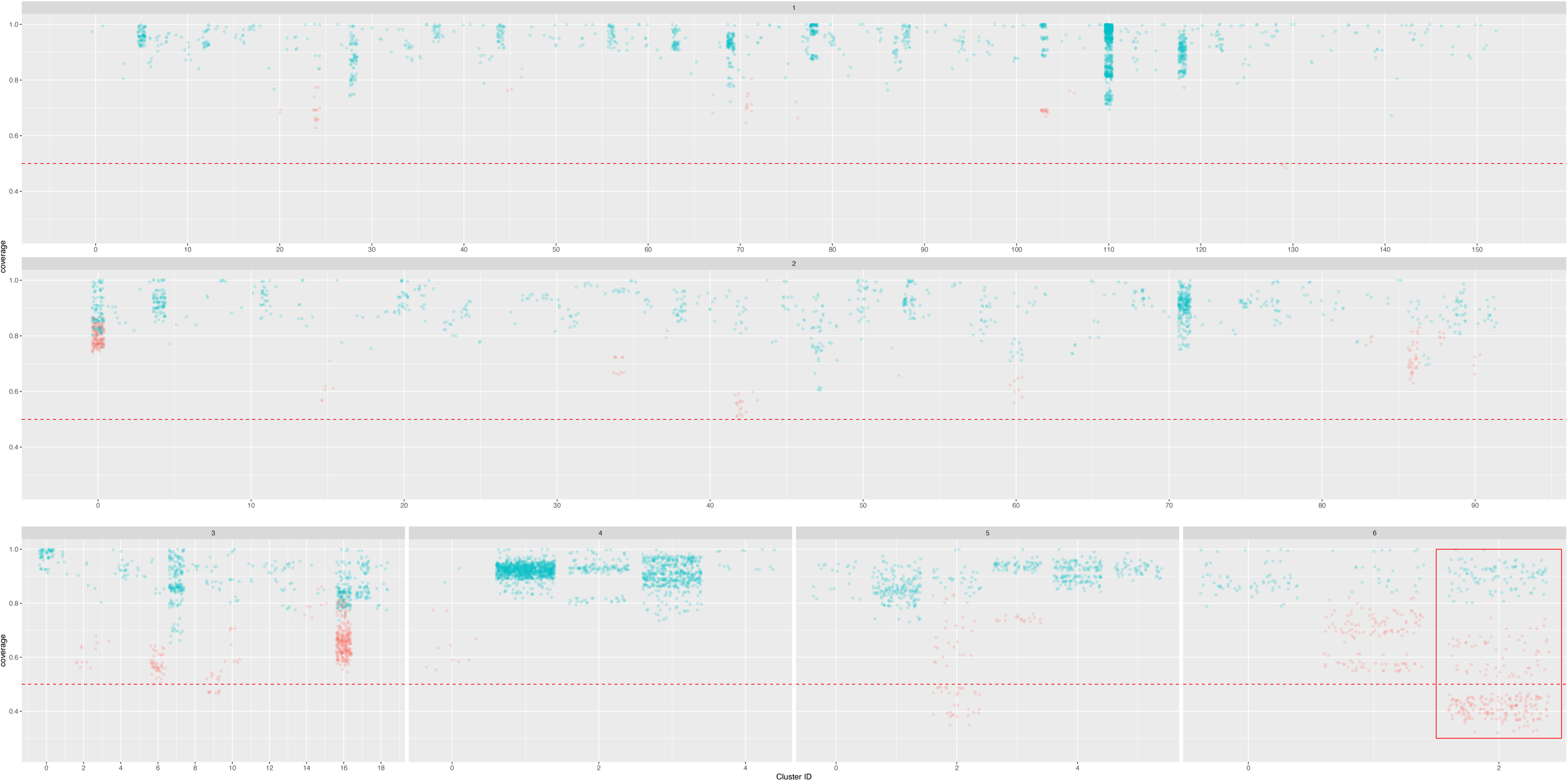
Scatterplots showing genome coverage for pairwise ANI comparisons for genomes sharing identical full-length and ambiguity symbol-free 16S sequences. The number of unique species names assigned per cluster is displayed at the top of each subgrouping, and the red horizontal line at 50% indicates the whole-genome genus threshold. Within-species pairwise comparisons (>!95% genome identity) are shown in blue, and between-species comparisons (<95% genome identity) are shown in red. Cluster uniting genomes with the lowest genome coverage is outlined in the red box.

**Fig. 7.**
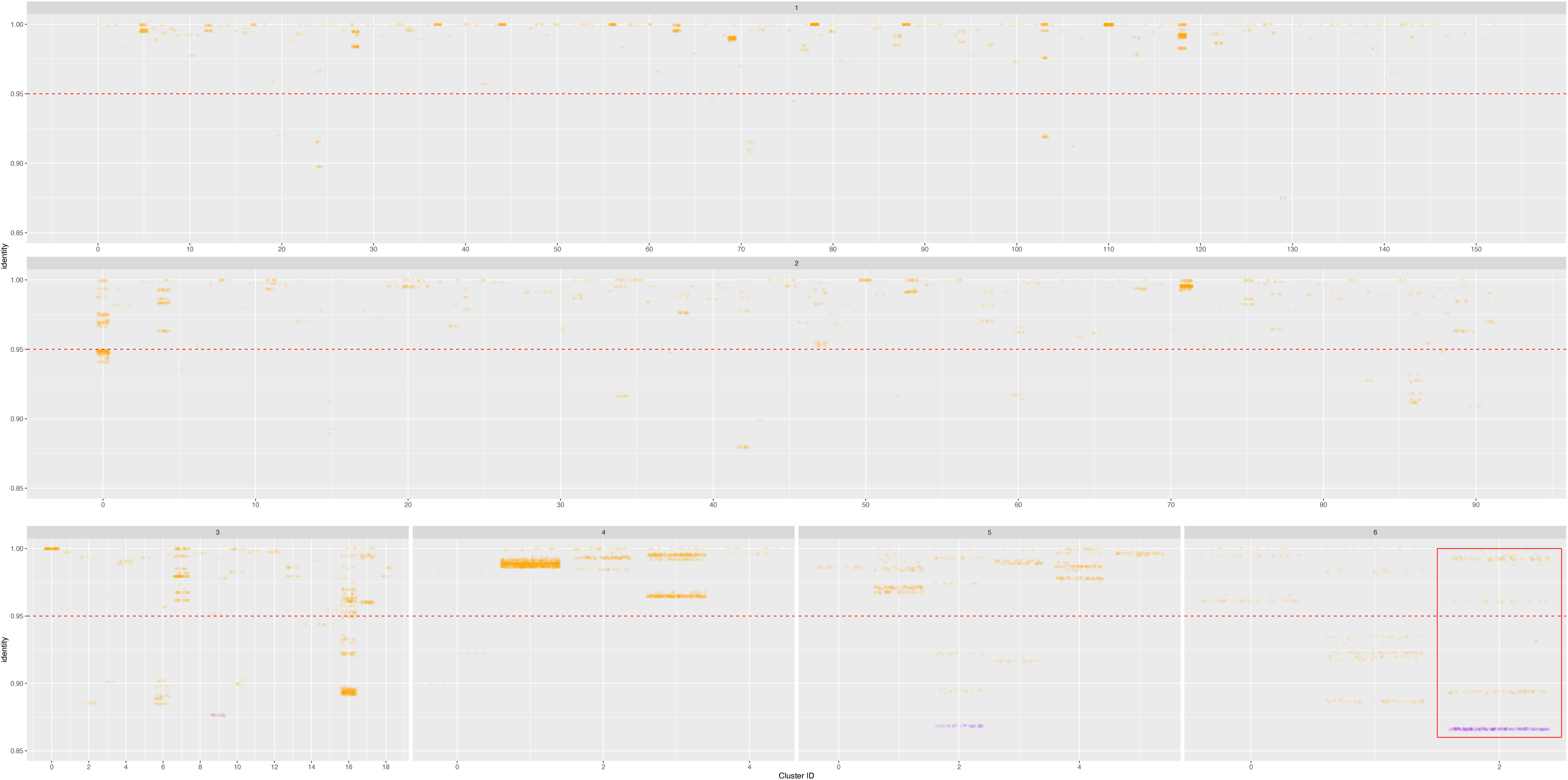
Scatterplots showing genome identity for pairwise ANI comparisons for genomes sharing identical full-length and ambiguity symbol free 16S sequences. The number of unique species names assigned per cluster are displayed at the top of each plot, and the red horizontal line at 95% indicates the whole-genome species threshold. Within-genus comparison (>!50% genome coverage) are shown in orange, and between-genus comparisons (<50% genome coverage) are shown in purple. Cluster uniting genomes with the lowest genome identity is outlined in the red box.

We find four (1.4%) clusters containing assemblies that share less than 50% genome coverage, some with as little as 32% coverage (Cluster 2, assigned six unique taxon names; **Fig. 6 – Red Box**). We find that, it is rare, but possible, for genomes from different candidate *Streptomyces* genera to share an identical 16S rRNA sequence. We also find that 36 (13%) of clusters include assemblies sharing less than 95% ANI (as low as 86% ANI in Cluster 2 with six unique taxon names; **Fig. 7**- red box). This observation is especially evident in, but not restricted to, clusters whose genomes have already been assigned distinct species names in NCBI. These results again show a one-to-many mapping between 16S sequence and *Streptomyces* genera and species as determined by whole-genome comparison, and that assignment of taxonomy based only on 16S rRNA sequences may be misleading.

However frequent the potential for misclassification, our data confirm that 98.6% of clusters comprise only representatives of a single genus, and 86% representatives of a single species, as determined by whole-genome comparison. Our *Streptomyces* genome sample is large but not exhaustive, so this may truly reflect that 16S rRNA sequences are often unique to a single species or genus. However, it remains possible that some of these 16S sequences may also be found in as yet unsequenced (or unreleased) assemblies with a different taxonomic classification. We note that some *Streptomyces* appear to be classified with more precision, potentially due to their industrial or medical importance: members of cluster 139, representing 27 genomes currently assigned to *Streptomyces clavuligerus*, share 96% coverage, and 100% identity. *S. clavuligerus* is an industrially important organism due to its ability to produce clavulanic acid [71]. By contrast, individual members of 125 (45%) clusters seem to have been assigned incorrectly to distinct species, as all members of the cluster share at least 50% genome coverage and 95% identity. Overall, our data show that, while many 16S rRNA sequences do resolve to single species level, there is not a one-to-one mapping between 16S and whole-genome taxonomy and, in general, 16S does not provide sufficient resolution to discriminate between species. We also conclude, on the basis of our observations, that extensive revision of the genus *Streptomyces* is required, in line with recent work based on a smaller dataset of 456 strains, suggesting that there are at least six validly-describable genera within the current single genus *Streptomyces* [31].

### Conclusions/Recommendations

The 16S rRNA gene is one of the most widely-used phylogenetic markers for studying microbial diversity [72] due to its ubiquitous distribution across all bacterial species [73]. 16S rRNA sequences are often clustered into OTUs at a canonical threshold of 97%, or as zOTUs/ASVs, as proxies for species [63]. With the large genomic datasets now available for *Streptomyces* species, we set out to address three questions: are 16S database sequences and taxonomic annotations of sufficient quality for use as a reference in taxonomic classification?; is 16S taxonomic classification a reasonable proxy for whole-genome classification in principle?; and what does the whole-genome classification tell us about *Streptomyces* taxonomy in general?

We collated *Streptomyces* 16S data from Greengenes, RDP, SILVA and NCBI databases, reducing a total dataset of ≈48,000 sequences to a high-quality dataset of ≈14,000 non-redundant full-length sequences, finding that around two-thirds of the public sequence data was either not of sufficient sequence or annotation quality for our analysis. We used Maximum Likelihood phylogeny of the high-quality dataset to obtain the largest 16S sequence tree published to date. Despite a relatively low level of bootstrap confidence, we found a clear partitioning of *Streptomyces* into three major clades, and that in some cases database taxonomic assignments placed the same species names at points in the topology inconsistent with a common lineage – in some cases placing species representatives into more than one major clade. By clustering zOTUs, we observe that in several cases, unique 16S sequences are associated with more than one current database taxonomic assignment, and that unique 16S sequences can be associated with more than one *Streptomyces* species, or even genus, when whole-genome approaches are used for classification. By surveying complete *Streptomyces* genomes, we show that single *Streptomyces* genomes may frequently contain multiple distinct 16S rRNA sequences. Taken together, these results demonstrate that there is a one-to-many relationship between 16S rRNA sequence and *Streptomyces* species, and a one-to-many relationship between *Streptomyces* species and 16S rRNA sequence. This calls into question the utility of 16S marker sequences for colony identification, metabarcoding, and environmental analyses in *Streptomyces*.

Finally, our whole-genome analyses of *Streptomyces* genomes imply that there is an implicit underestimation of genome-level diversity in the *Streptomyces* taxonomy. We find multiple groups of genomes, currently united in the *Streptomyces* genus, that are dissimilar at the same level as genomes accepted to be distinct genera in other lineages of bacteria. We additionally find that, using a heuristic threshold for ANIm-based species assignment, there are grounds for reassignment of many genomes, including the unification and creation of new, species-level groupings. Our data suggest that revision of the genus *Streptomyces*, using whole-genome methods, is essential and overdue.

## Author Contributions

Conceptualization: A.B.K, P.A.H and L.P

Data curation: A.B.K and L.P

Formal analysis: A.B.K

Funding acquisition: P.A.H and L.P

Methodology: A.B.K, P.A.H and L.P

Project administration: A.B.K and L.P

Supervision: P.A.H and L.P

Writing – original draft: A.B.K, P.A.H and L.P

Writing – review and editing: A.B.K, P.A.H and L.P

## Conflicts of interest

The authors declare that there are no conflicts of interests.

## Funding Information

This research was supported by a grant from the University of Strathclyde Research Excellence Award. LP would like to acknowledge funding from BBSRC (BB/V010417/1). PAH would like to acknowledge funding from BBSRC (BB/T001038/1 and BB/T004126/1) and the Royal Academy of Engineering Research Chair Scheme for long term personal research support (RCSRF2021\11\15).

## Ethical approval

Not applicable

## Consent for publication

Not applicable

## Abbreviations

AAI: average amino acid identity
AMR: antimicrobial resistance
ANI: average nucleotide identity
ASV: Amplicon Sequence Variants
dDDH: digital DNA–DNA hybridization
LPSN: List of Prokaryotic names with Standing in Nomenclature
ML: maximum likelihood
MLSA: Multi Locus Sequence Analysis
NCBI: National Center for Biotechnology Information
OTUs: opertaional taxonomic unit
RDP: Ribosomal Database Project
TBE: Transfer Bootstrap Expectation
zOTUs: zero-radius operational taxonomic units

## Acknowledgements

We acknowledge the University of Strathclyde Research Excellence Award for funding our project and the ARCHIE-West computer cluster for computational resources and support.

## Funding Statement

The funders had no role in the design or decision to submit the work for publication

## Notes

https://doi.org/10.5281/zenodo.8223787

### Competing Interest Statement

The authors have declared no competing interest.

